# Serial fermentation in milk generates functionally diverse community lineages with different degrees of structure stabilization

**DOI:** 10.1101/2024.04.01.587544

**Authors:** Chloé Gapp, Alexis Dijamentiuk, Cécile Mangavel, Cécile Callon, Sébastien Theil, Anne-Marie Revol-Junelles, Christophe Chassard, Frédéric Borges

**Affiliations:** Université de Lorraine, LIBio, Nancy, France; Université Clermont Auvergne, INRAE, VetAgro Sup, UMR 0545 Fromage, Aurillac, France

**Keywords:** microbiome engineering, community structure, ecological trajectory, serial propagation, backslopping, lactic acid bacteria, raw milk, fermentation

## Abstract

Microbial communities offer considerable potential for tackling environmental challenges by improving the functioning of ecosystems. Top-down community engineering is a promising strategy that could be used to obtain communities of desired function. However, the ecological factors that control the balance between community shaping and propagation are not well understood. Dairy backslopping can be used as a model engineering approach to investigate the dynamics of communities during serial propagations. In this study, 26 raw milk samples were used to generate lineages of 6 communities obtained by serial propagation. Bacterial community structures were analyzed by metabarcoding and acidification was recorded by pH monitoring. The results revealed that different types of community lineages could be obtained in terms of taxonomic composition and dynamics. Five lineages reached a repeatable community structure in a few propagation steps, with little variation between the final generations, giving rise to stable acidification kinetics. Moreover, these stabilized communities presented a high inter-lineage variability of community structures as well as diverse acidification properties. Besides, the other lineages were characterized by different levels of dynamics leading to parallel or divergent trajectories. The functional properties and dynamics of the communities were mainly related to the relative abundance and the taxonomic composition of lactic acid bacteria within the communities. These findings highlight that short-term schemes of serial fermentation can produce communities with a wide range of dynamics and that the balance between community shaping and propagation is intimately linked to community structure.

**Importance:** Microbiome applications require approaches for shaping and propagating microbial communities. Shaping allows the selection of communities with desired taxonomic and functional properties, while propagation allows the production of the biomass required to inoculate the engineered communities in the target ecosystem. In top-down community engineering, where communities are obtained from a pool of mixed microorganisms by acting on environmental variables, a major challenge is to master the balance between shaping and propagation. However, the ecological factors that favor high dynamics of community structure and, conversely, those that favor stability during propagation are not well understood. In this work, short-term dairy blacksloping was used to investigate the key role of the taxonomic composition and structure of bacterial communities on their dynamics. The results obtained open up interesting prospects for the biotechnological use of microbiomes, particularly in the field of dairy fermentation, to diversify approaches for injecting microbial biodiversity into cheesemaking processes.

## Introduction

Microbiome engineering is an important source of innovations that could contribute to a more sustainable future and could be used to address a wide range of environmental, food, and health issues (1). It is defined as approaches dedicated to improving the function of an ecosystem by manipulating the composition of microbes (2). Despite this high potential, the great complexity of microbiomes is a major barrier to the emergence of such breakthrough innovations, as most engineering approaches in microbiology are adapted to simple microbiological systems. Therefore, efforts are being made using model microbiomes to develop engineering approaches based on the understanding of ecosystem functioning in a controlled environment (1). In this context, food fermentations are interesting models because these ecosystems are relatively simple, easy to control and, due to their ancestral anthropic origin, can be a rich source of inspiration for microbiome engineering (3).

During cheesemaking, microorganisms play a major role at all stages of processing including acidification, alkalinization, and ripening (4). They participate in texture changes, aroma development, and can also preserve cheese from pathogen colonization (5, 6). In the acidification step, lactic acid bacteria (LAB) are mainly responsible for the production of lactic acid using lactose from the milk (7). LAB are Gram-positive, non-spore forming, catalase-negative, and tolerant to acidic pH. LAB encompass the genera from the order *Lactobacillales* including *Enterococcus, Lactobacillus sensu lato*, *Lactococcus*, *Leuconostoc*, *Oenococcus, Pediococcus*, *Streptococcus*, and *Weissella* (8, 9). In cheese manufacturing, the LAB used as starters for acidification are *Lactobacillus sensu lato*, *Lactococcus* and *Streptococcus* (10). However, these starters are characterised by a low diversity and, despite their widespread use in industry, there are strong expectations in the cheese sector for innovations using microbiome engineering approaches.

There are two main approaches to community design: bottom-up and top-down (11). The bottom-up method consists in assembling individual microorganisms of interest into communities. The top-down approach consists in applying environmental pressures to select adapted communities, allowing propagation and shaping of communities. One of the main challenges of top-down community engineering is to master the balance between conservative propagation and shaping of communities. In this regard, two aspects are considered: the structure of the community and its function. The engineering processes can indeed be designed in order to shape, or preserve, the community structure or its function, depending on the purpose. A number of community propagation technologies have been proposed, enabling these communities to be propagated with varying degrees of effect on their characteristics (12–14). Among these, sequential propagation is particularly interesting because it has been used empirically for a very long time in traditional food fermentation. This inoculation technique called backslopping consists in using a small fraction of a previous production to inoculate the new one. Backslopping can be considered as a top-down microbiome engineering approach as it allows the selection of communities increasingly adapted to technological pressures (milk, temperature, salt, pH) while assuring desired functions (11) such as acidification. Omics approaches have been used to unravel key features of microbial community dynamics during backslopping, and the consequences for community function (15–20). Long-term backslopping leads to communities with low species richness and high intraspecies diversity (21, 22). Although the composition of these communities varies, their function is highly similar, revealing a functional redundancy conferring functional stability on the communities (21, 22). However, these studies were carried out on communities resulting from a long backslopping process involving a large number of fermentation batches, and therefore very little is known about how these communities evolve during the initial stages of backslopping. A recent study has shown that it is possible to obtain communities whose structure is stabilised on an interspecific scale (23). However, this study used three samples of raw milk and the propagations were carried out in laboratory medium, which does not allow us to take into account the microbial diversity of raw milk and the behaviour of the communities when the propagations are carried out in milk.

The aim of this study was to investigate the dynamics of bacterial communities during the early stages of a backslopping process, and to analyze the relationship between the dynamics of community structures and their acidification properties. We set up an experiment based on serial fermentation starting from raw milk samples collected from 26 different farms of the same PDO geographical area in the north east of France. Each milk sample was serially fermented 6 times in mesophilic conditions. Acidification kinetics were recorded by pH monitoring during each fermentation and bacterial communities were analyzed by metabarcoding.

## Material and methods

### Sample collection

Milk samples, called M1 to M26, were collected from the tanks of 26 farms in the north east of France (Grand Est region). Approximately 500 mL was taken directly from the tanks for each sample, put in a sterile container and kept at a low temperature until arrival at the laboratory. Samples were stored at 4 °C overnight. Then 300 mL was used for centrifugal concentration for DNA extractions and 25 mL was stored at −80 °C until serial fermentation.

### Raw milk concentration

To concentrate raw milk for the metabarcoding analysis, 300 mL of each milk sample was centrifuged for 30 minutes at 5,300 g at 4 °C. Liquid and cream were discarded and the pellets were resuspended in 1.5 mL of phosphate-buffered saline. Then, the samples were put in 2 mL tubes and centrifuged for 5 min at 13,000 g at 4 °C. Supernatants were discarded and the pellets were stored at −80 °C until DNA extraction.

### Serial fermentation

Tubes containing 25 mL of frozen milk were thawed and placed in a water bath at 34 °C for 24 h. Then, 250 µL of each fermented sample was inoculated in 25 mL of fresh UHT milk (Marque Repère, E. Leclerc, Ivry-sur-Seine, France) and incubated for 18 h at 34 °C. This step was repeated 5 times to get 6 fermentations in total for each milk sample (Figure 1). During each fermentation, the pH was monitored and at the end of each fermentation samples were collected, placed in 2 mL tubes, and stored at −80 °C until DNA extraction.

**Figure 1.**
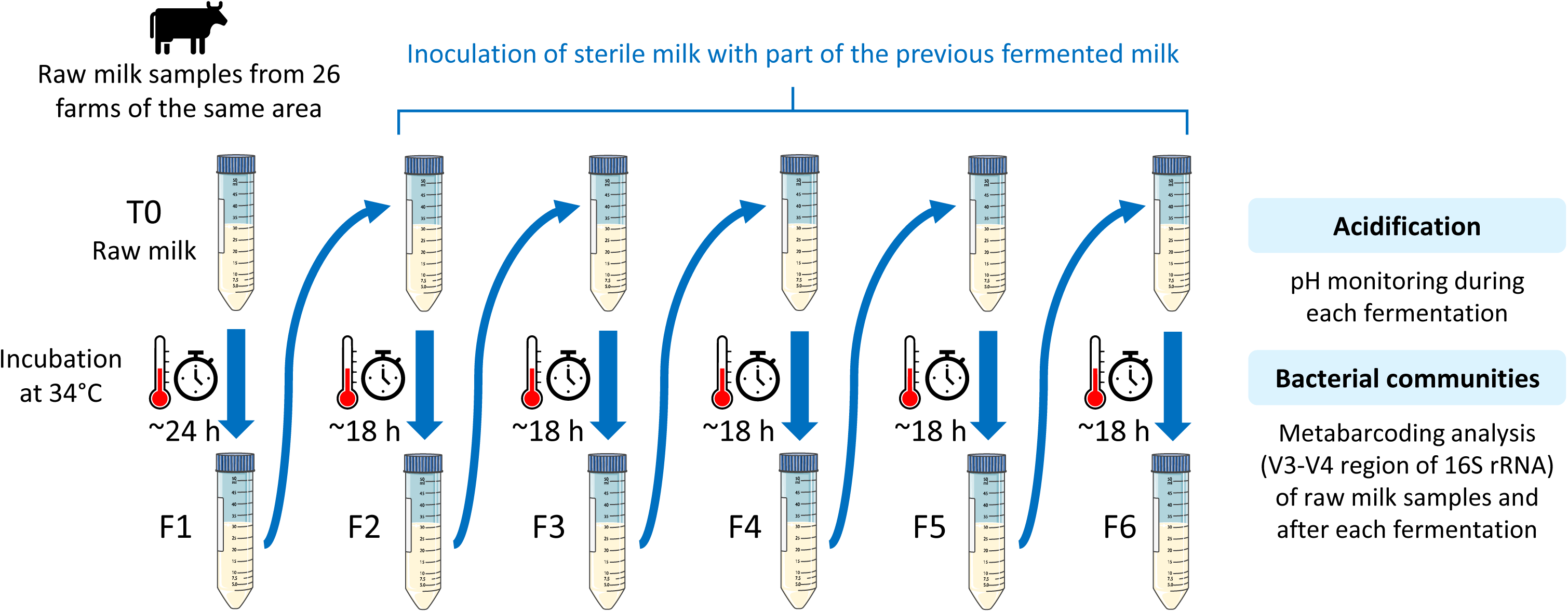
Diagram summarizing the serial fermentation protocol. Raw milk samples collected from 26 farms were incubated for 24 h at 34 °C to allow spontaneous fermentation. Then, sterile milk was inoculated at 1:100 with the fermented milk and incubated for 18 h at 34 °C and this process was repeated 5 times to obtain 6 fermented samples for each starting raw milk.

### Acidification kinetics

Acidification was recorded by pH monitoring during each fermentation. Before each pH time course, pH sensors (VWR pHenomenal 111, VWR international, Radnor, PA, USA) were cleaned with deionised water and sterilised by immersion in 70% ethanol, then rinsed by immersion in sterile deionised water. The pH was measured every 5 minutes using a multi-channel pH meter (Consort datalogger D291, Consort bvba, Turnhout, Belgium).

### Acidification parameter analysis

Acidification parameters (lag time, maximum acidification rate and final pH) were extracted from each curve using a custom R script (version 4.3.1, R Core Team, 2023).

Maximum acidification rate (MAR) was the lowest slope coefficient among the linear regressions obtained on a 20-point sliding window. The resulting linear model was used to calculate the intersection point with a line of null slope and intersecting y axis at the first pH value of each time point. The time point corresponding to the intersection between this line and the linear model for the MAR was used as an estimation of the lag time (corresponding to the time before the pH drop). For the final pH, a linear model was built with the last 20 points of the curves and used to predict the final pH at 18 h (for F2 to F6) or 24 h (for F1).

### DNA extraction

DNA was extracted from the raw milk pellets and the fermented samples using a FastDNA Spin Kit for Soil (MP Biomedicals, Illkirch-Graffenstaden, France) with bead beating, following the manufacturer’s recommendations. DNA was stored at −20 °C. Contamination controls were performed following the same protocol without DNA, every time extractions were performed.

### PCR amplification

A nested PCR was performed for raw milk samples. First, the 16S rRNA gene (∼1450 bp) was amplified using Taq polymerase (Appligene, Illkirch-Graffenstaden, France), and the universal bacterial primers W02 (5’-GNTACCTTGTTACGACTT-3’) and W18 (5’-GAGTTTGATCMTGGCTCAG-3’) as described previously (24, 25). The PCR program was initiated with incubation at 94 °C for 3 min, followed by 17 cycles of 94 °C for 30 s, 50 °C for 30 s and 72 °C for 1 min 30 s, followed by a final extension at 72 °C for 10 min. Then, the product obtained with this first PCR was used as a matrix for the next PCR, targeting the variable region V3-V4 of the 16S rRNA gene (∼510 bp) using MTP taq polymerase (Sigma, Saint-Louis, MO, USA) and the primers MSQ-16SV3F (5′-TACGGRAGGCWGCAG-3′) and PCR1R_460 (5′-TTACCAGGGTATCTAATCCT-3′), as described previously (26). The PCR program was initiated with incubation at 94 °C for 1 min, followed by 30 cycles of 94 °C for 1 min, 65 °C for 1 min, and 72 °C for 1 min, followed by a final extension at 72 °C for 10 min. For fermented samples, only the second PCR for V3-V4 amplification was performed using the extracted DNA as matrix.

### Sequencing and data analysis

The 16S amplicon sequencing was performed by the GeT platform (GenoToul, Toulouse, France) using the Illumina Miseq technology, for the 26 raw milk samples and every fermented sample (182 total), as well as the contamination controls for the DNA extraction. Data were analyzed using the rANOMALY workflow (27), to produce amplicon sequence variants (ASV) via the ‘dada2’ package (28), filter contaminations and extract the abundance table. Then, taxonomic affiliation was made using FROGS tools (29) on the Migale Galaxy server (30) with the 16S EZBioCloud 52018 database (31).

Further analysis was performed using R custom scripts (R version 4.3.1) (32). Alpha and beta diversity analyses were conducted using tools from the ‘vegan’ package (33).

### Statistical analysis

Statistical tests were performed with R version 4.3.1 (32) using tools from the ‘rstatix’ (34), ‘biostat’ (35) and ‘segmented’ (36) packages.

Results were expressed as means ± standard error of the mean (SEM). Pairwise comparisons were performed with the Tukey honestly significant difference (HSD) test following analysis of variance (ANOVA) or the Wilcoxon rank sum exact test with the Bonferroni *p* value adjustment method following a Kruskal-Wallis rank sum test, when the data did not meet the application conditions. Variance comparisons were carried out using an F test. Correlation tests were carried out using Pearson’s product-moment correlation test. Simple linear regressions were performed to further investigate the relationship between quantitative variables. In the figures, different superscript letters indicate a significant difference (*p* < 0.05).

## Results

### Acidification dynamics during serial fermentation

During each fermentation step, the pH was monitored over time (Figure S1). Three parameters of interest were extracted from each pH time course: the time at which pH starts to drop (lag time), the maximum acidification rate (MAR), and the pH at the end of the fermentation (final pH).

The analysis of the parameters showed that the lag time was always longer for the first fermentation compared to the following steps (Figure S2A, *p* = 6.0 × 10^-14^, Wilcoxon rank sum exact test - WRSET). It took 14.3 hours on average before the pH drop occurred in the F1 and from 2.1 to 2.7 h for the following steps. The speed of pH drop was lower for fermentation F1 (−0.27 pH units per hour) compared to the other fermentations (from −0.36 to −0.39, Figure S2B, *p* < 9.0 × 10^-4^, WRSET). These results show that the first fermentation was markedly different from the others.

When the fermentations F2 to F6 were considered, visual inspection delineated three groups called GP1, GP2, and GP3 (Figure S1, examples shown in Figure 2A) among the 26 lineages. GP1 was characterized by highly reproducible acidification profiles from F2 to F6 (5/26 samples, Figure S1), whereas in GP2 the acidification curves showed small observable variations from F2 to F6 (9/26 samples, Figure S1). In GP3, in contrast to the other groups, the change in acidification was more variable, with intersecting curves (12/26 samples, Figure S1).

**Figure 2.**
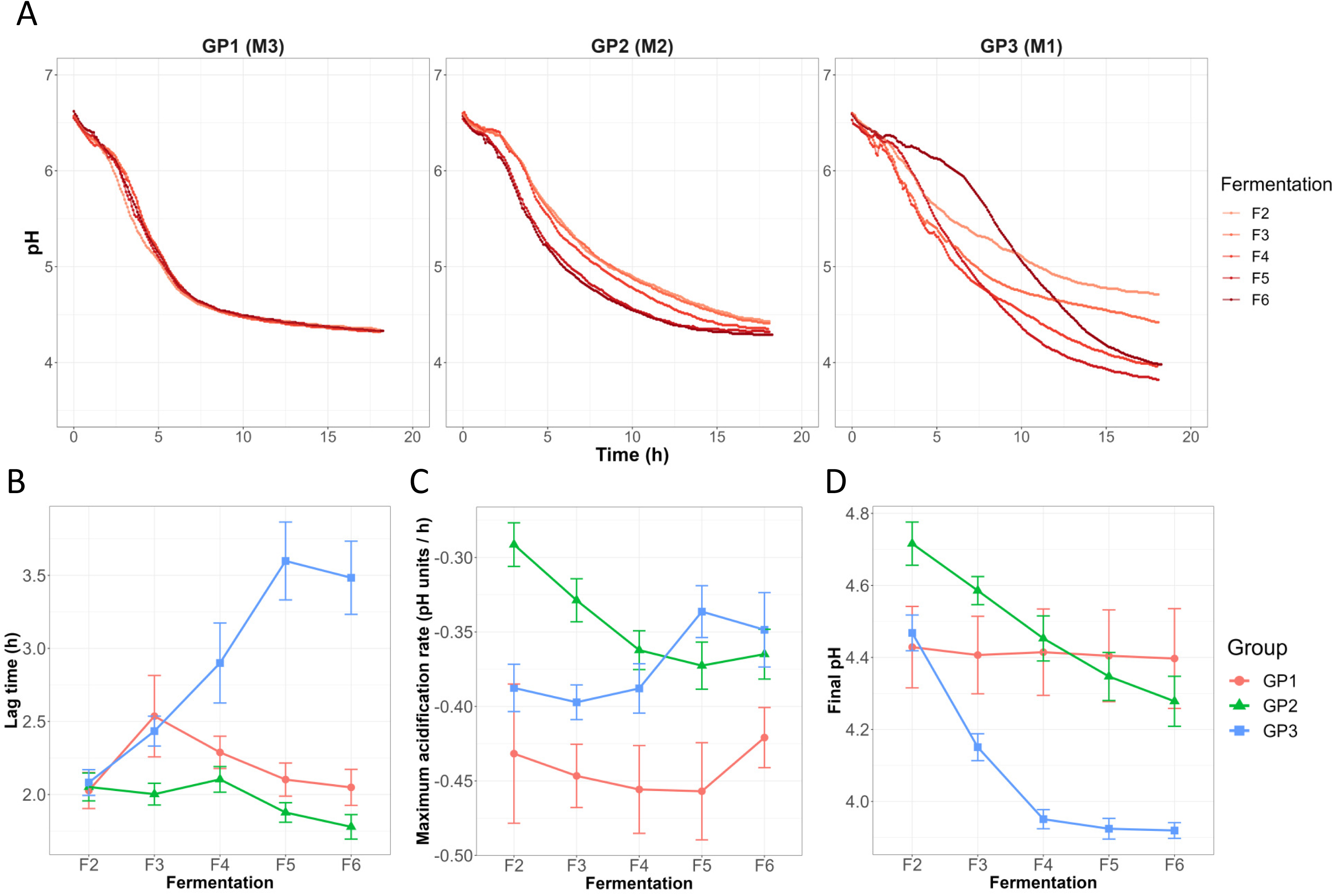
Analysis of acidification kinetics. (A) Examples of acidification kinetics showing the different patterns in the three groups: M3, M2, and M1 for GP1, GP2, and GP3, respectively. The acidification parameters (means and standard errors) lag time, MAR and the final pH are represented in panels B, C, and D, respectively.

In GP1, the lag time, the MAR, and the final pH did not significantly change over the course of fermentation (Figures 2B, 2C, 2D, *p* > 0.05 Pearson’s product-moment correlation – PPMC). Interestingly, even if the final pH in GP1 was stable from F2 to F6, it was significantly different among the lineages: the final pH was 4.13 for M15 on average, 4.33 for M3 and M26, 4.40 for M18, and 4.86 for M19 (all lineages were significantly different - *p* < 0.022 - except lineages M3 and M26 according to the Tukey HSD test following ANOVA). By contrast, for GP2 the lag time decreased (Figure 2B, r = −0.36, *p* = 1.4 × 10^-2^, PPMC), the speed of pH drop increased (Figure 2C, r = −0.52, *p* = 2.6 × 10^-4^, PPMC), and the final pH decreased (Figure 2D, r = −0.67, *p* = 3.7 × 10^-7^, PPMC). A significant change in the acidification parameters was also observed for GP3: the lag time greatly increased (Figure 2B, r = 0.61, *p* = 2.1 × 10^-7^, PPMC), the speed of pH drop decreased (Figure 2C, r = 0.31, *p* = 1.7 × 10^-2^, PPMC), and the final pH significantly decreased from F2 to F4 (Figure 2D, r = −0.85, *p* = 6.9 × 10^-11^, PPMC), and then remained stable from F4 to F6 (r = −0.15, *p* = 0.39, PPMC). Overall, these results show that three groups can be delineated according to their acidification patterns from the second fermentation step onwards. The group GP1 was characterized by stable acidification parameters, but contained lineages characterized by different final pH values. In contrast, the other two groups GP2 and GP3 were characterized by different changing acidification kinetics during the serial propagations.

### Bacterial community dynamics during serial fermentation

Overall, bacterial community structures of the raw milk samples and of serial fermentation revealed a high variability in terms of both taxonomic composition and dynamics (Figure 3). The majority of initial raw milk samples (17/26) contained more than 50% relative abundance of the *Pseudomonas* genus (with species *P. yamanorum* for 16 samples and *P. gessardii* for 1 sample, T0 Figure 3). One sample (M6) was dominated by *Streptococcus porcinus* whereas other samples were more diversified (M18, M21, M20, M22, M4, M17, M19, M16, with a richness index of more than 75 and a Shannon index above 2.5, Figures S3A, S3C). No significant differences in richness and Shannon index were observed between the starting raw milk samples of the 3 groups (Figures S3B, S3D).

**Figure 3.**
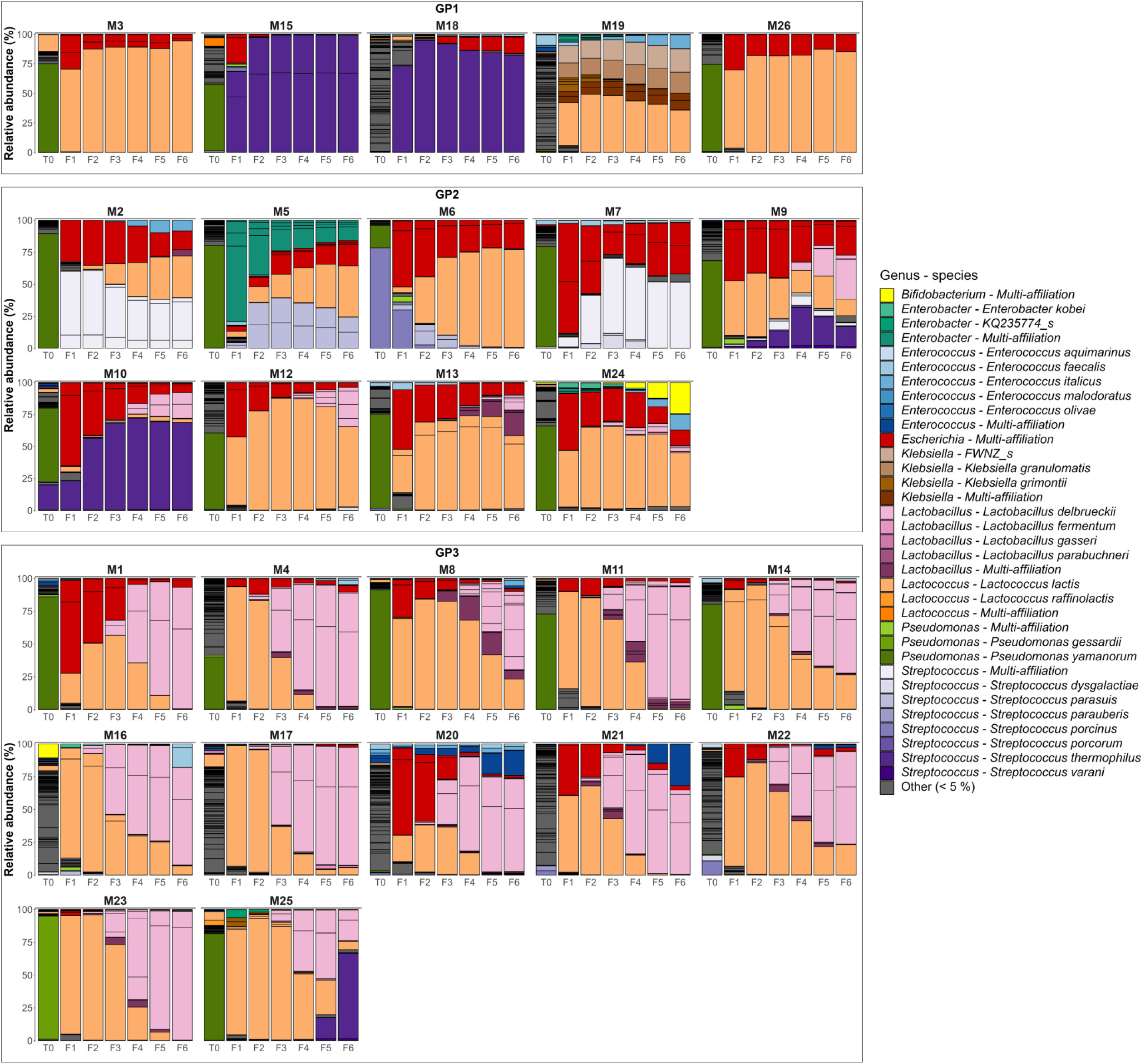
Bacterial composition of the community lineages at the genus or species level. T0: raw milk samples, F1 to F6: fermented milk samples. Genera representing less than 5% of the total relative abundance for each fermentation series have been aggregated into the class “Other” (grey). The barplot series are presented according to groups GP1, GP2, and GP3.

After one fermentation (F1), the richness markedly decreased for all lineages from 60.9 ± 3.9 in T0 to less than 23.0 (Figure S4A, *p* < 1.6 × 10^-8^, WRSET). The richness in the last fermentation step was 13.2 ± 0.7. The first fermentation also exhibited a decreased Shannon index (from 1.9 ± 0.2 to 1.2 ± 0.1), although less noticeably (Figure S4B, only the comparison of T0 with F2 was significant with *p* = 1.3 × 10^-2^, WRSET, otherwise *p* > 0.19). The poor significance is likely due to a higher variability of the Shannon index between raw milk samples compared to propagated communities (Figure S4B, *p* < 2.2 × 10^-16^, F test to compare the variances between T0 and fermented samples). These results showed that serial fermentation induces a decrease in bacterial alpha diversity.

Among the F1 communities, 17/26 were dominated by ASV of the phylum *Bacillota* (Figure S5), and more specifically by LAB ASV from the genera *Lactococcus* or *Streptococcus* (with species *L. lactis* and *S. thermophilus*, respectively, Figure 3). The other remaining F1 communities were dominated by Gram-negative bacteria of the phylum *Pseudomonadota* (*Escherichia* genus in 7 F1 communities, *Klebsiella* genus for one and *Enterobacter* genus for another one). In all cases, the communities ended up being dominated by *Bacillota* from step F3. However, the *Pseudomonadota* remained present in a large proportion until fermentation F6 in several fermented milk samples (*e.g.* M6, M7, M9). Interestingly, the abundance of *Lactobacillus* genus *sensu lato* was very low in F1 (< 1%) and started to increase to at least 25% (up to 63%) in 9 F3 communities. They ended up being dominant (> 50%) at F6 for 11 lineages where the main species was *L. delbrueckii*.

The bacterial communities from GP1 were characterized by minor changes in community structure during serial fermentation (Figure 3). *Lactococcus* (M3, M26) or *Streptococcus* ASV (M15, M18) was dominant in GP1 (> 75% relative abundance), except for M19 in which proportions of *Lactococcus* and *Klebsiella* were similar during the fermentation steps, with *Klebsiella* even becoming dominant in the last steps (Figure 3). The GP2 communities showed more changes than GP1 communities and were dominated by multiple LAB (*Lactococcus*, *Streptococcus*, *Lactobacillus* and *Enterococcus,* Figure 3). GP3 showed broader changes in the communities: while *Lactococcus* ASV were dominant during the first propagation steps, they were subsequently replaced by *Lactobacillus* ASV *sensu lato* during the course of serial fermentation (Figure 3). The relative abundance of *Lactobacillus sensu lato* was significantly higher in GP3 compared to GP2 (*p =* 6.2 × 10^-10^, WRSET), and in GP2 compared to GP1 (*p =* 2.2 × 10^-4^, WRSET). These results revealed that community structures from GP1, GP2 and GP3 exhibit different patterns of change.

The analysis of richness and Shannon index showed that GP1 was less diversified than GP2 and GP3 (Figures S6B, S6E). In addition, the groups appeared to differ in the range of variation of these metrics over the course of serial fermentation (Figures S6A, S6D). Therefore, the sums of the differences between adjacent fermentation steps (n+1 and n) were calculated to report specifically on these variations in alpha diversity. The results show that Shannon diversity fluctuated significantly more in GP3 compared to GP1 and GP2 (Figure S6F, *p* < 0.0098, WRSET). On the other hand, no significant differences were observed between the groups, regarding the variation in richness (Figure S6C).

Bray-Curtis dissimilarity was calculated for each fermentation series, between the fermentation steps n and n+1, from F2 to F6. GP1 showed a stable Bray-Curtis dissimilarity level between each adjacent step (*p* = 0.90, PPMC), and was relatively low (0.03 ± 0.01, Figure 4A). The dissimilarity indices in GP2 were higher (0.13 ± 0.01, *p* = 1.2 × 10^-8^, WRSET) than in GP1 and were also stable (Figure 4A, *p* = 0.18, PPMC). For GP3, Bray-Curtis dissimilarity significantly decreased (r = −0.47, *p* = 7.2 × 10^-4^, PPMC): from 0.31 ± 0.05 between F2-F3 to 0.14 ± 0.04 between F5-F6. Overall, the sum of Bray-Curtis dissimilarity between adjacent steps increased significantly from GP1 to GP3 (Figure 4B, *p* < 0.001, Tukey HSD test following ANOVA). These results show that the three groups GP1, GP2, and GP3, are characterized by different levels of community structure dynamics during the course of serial fermentation.

**Figure 4.**
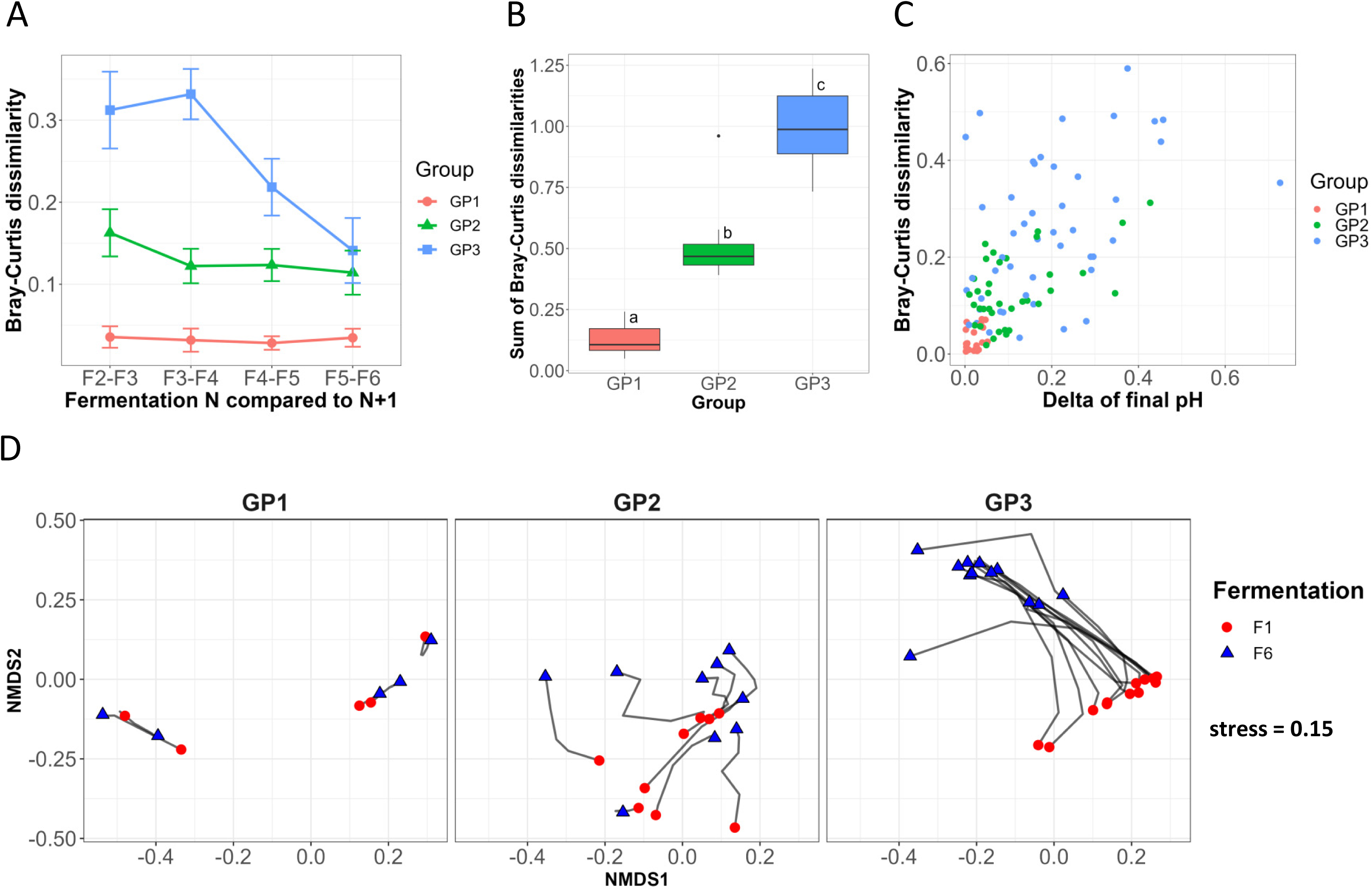
Beta diversity analysis. (A) Mean Bray-Curtis dissimilarities between adjacent fermentation steps from F2 to F6. Error bars represent the standard errors of the means. (B) Sums of Bray-Curtis dissimilarities between adjacent steps for each group of communities. Different superscript letters indicate a significant difference (*p* < 0.05; Pairwise comparisons using the Wilcoxon rank sum exact test with the Bonferroni adjustment method for the *p* value). (C) Bray-Curtis dissimilarities and delta of final pH between adjacent steps from F2 to F6. (D) Non-metric multi-dimensional scaling ordination representing ecological trajectories for each group.

The non-metric multidimensional scaling (NMDS) ordination based on Bray-Curtis dissimilarity revealed that GP1 community lineages were stable but segregated, showing a high inter-lineage variability (Figure 4D). GP2 lineages showed longer ecological trajectories than GP1 and their directions were divergent. In GP3, communities were highly dynamic and were characterized by longer trajectories. Interestingly, the direction was parallel for the majority of the GP3 lineages, showing that the change in taxonomic composition was similar in those communities compared to the other groups. These results show that the groups are characterized by different typologies of lineage trajectories and segregation.

A possible correlation was investigated between the acidification properties of the communities and Bray-Curtis dissimilarity. For this purpose, the difference in pH metrics between adjacent steps from F2 to F6 was calculated for each lineage. A significant correlation was highlighted between the delta of the final pH and Bray-Curtis dissimilarity: the higher the pH variation, the greater the Bray-Curtis dissimilarity (Figure 4C, r = 0.62, *p* = 2.5 × 10^-12^, PPMC). No significant correlation was obtained with the delta of lag or MAR. This result reveals that the stability of community structures is related to functional stability.

### Relationship between acidification properties and community structure

LAB are known to be responsible for acidification by producing lactic acid from lactose in milk. As they were highly abundant in the communities, we focused on the impact of LAB on acidification properties.

The proportion of LAB only slightly impacted the acidification lag time when they represented less than 88% of relative abundance within the communities (*p* = 0.10 for the linear model, Figure 5A). However, when the communities contained more than 88% LAB, the lag time significantly increased (slope = 0.090, *p* = 4.4 × 10^-3^ for the linear model). This increase in lag time mostly concerned communities dominated by *Lactobacillus sensu lato* ASV (3.19 ± 0.16 h on average, Figure 5D) compared to the other communities (from 1.95 ± 0.04 to 2.14 ± 0.05 h, *p* < 2.3 × 10^-5^, WRSET).

**Figure 5.**
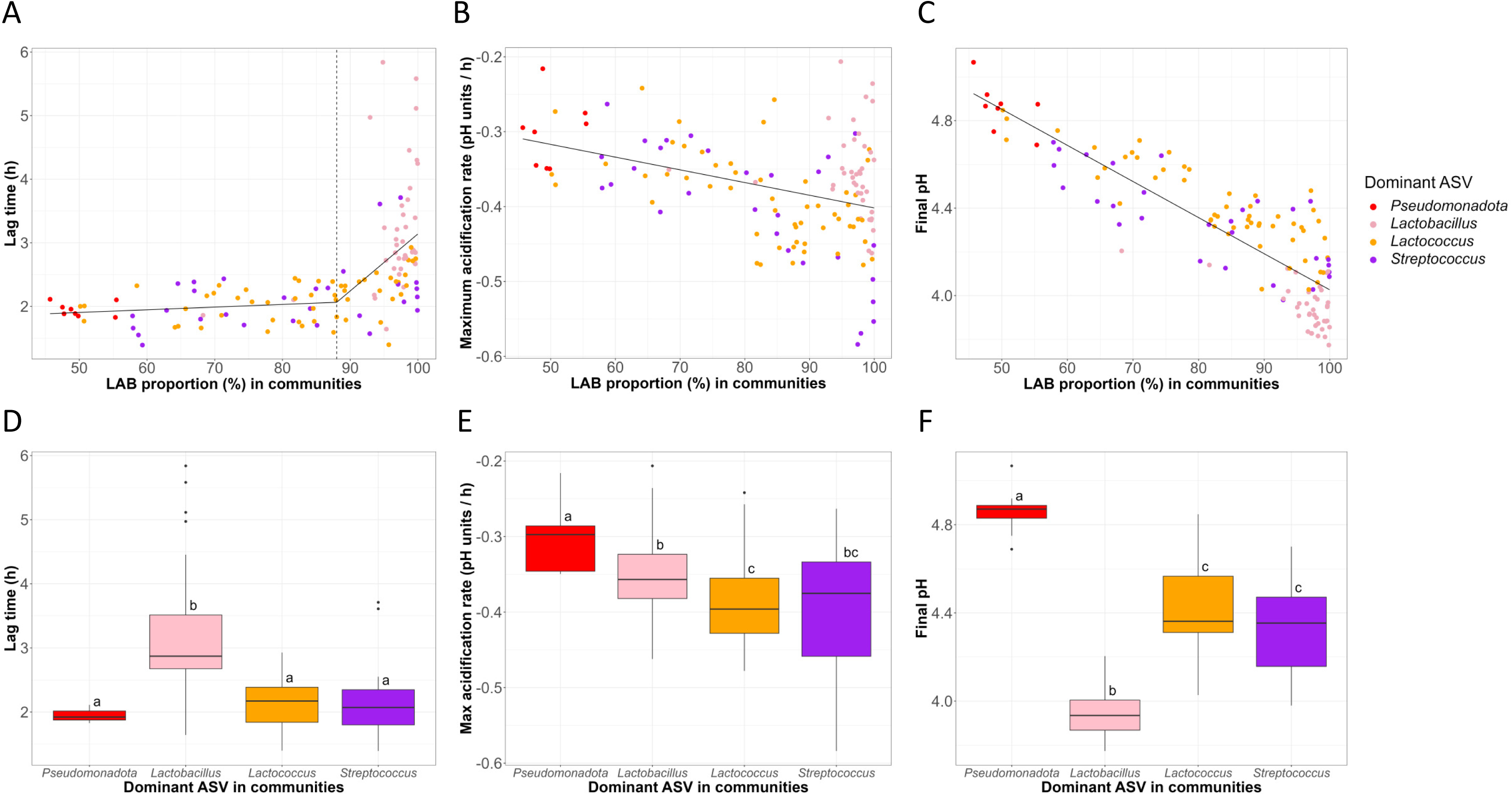
Relationship between the acidification parameters and the LAB proportion and composition. The lag time, the MAR, and the final pH as a function of the proportion of LAB in communities (A, B, C, respectively), or the dominant taxa in communities (D, E, F, respectively). Different superscript letters indicate a significant difference (*p* < 0.05; Pairwise comparisons using the Wilcoxon rank sum exact test with the Bonferroni adjustment method for the *p* value).

In addition, when the relative abundance of ASV corresponding to LAB increased, the speed of pH drop increased and the final pH decreased (*p* = 5.8 × 10^-6^ for MAR, *p* < 2.2 × 10^-16^ for the final pH, Figures 5B, 5C). Accordingly, when the communities were dominated by *Pseudomonadota* ASV, the decrease of pH was slower compared to communities dominated by LAB (Figure 5E). The MAR also depended on the LAB genus: the decrease in pH was significantly slower when the communities were dominated by *Lactobacillus sensu lato* ASV (−0.35 ± 0.01 pH units per hour) compared to *Lactococcus* (−0.39 ± 0.01 pH units per hour, *p* = 3.5 × 10^-2^, WRSET, Figure 5E). On the other hand, a high variability in the speed of pH drop was observed when the communities were dominated by *Streptococcus* ASV (Figure 5E). The final pH of fermentation was strongly related to the dominant ASV, and was the highest for communities dominated by *Pseudomonadota* ASV (4.86 ± 0.04, *p* < 7.9 × 10^-5^, WRSET, Figure 5F). Conversely, the final pH was the lowest when *Lactobacillus sensu lato* ASV (3.95 ± 0.02) were dominant (*p* < 3.8 × 10^-8^, WRSET, Figure 5F). The communities dominated by the *Lactococcus* or *Streptococcus* ASV were characterized by intermediate final pH (4.41 ± 0.03 and 4.34 ± 0.04, respectively, Figure 5F).

These results show that the acidification properties are linked to the proportion of LAB in the community and more specifically to the dominant LAB genus.

## Discussion

We have investigated the changes in bacterial community structures and their acidification properties during experimental backslopping in milk. To do so, 26 raw milk samples were incubated and the resulting communities were serially propagated in sterile milk. Communities were analyzed by metabarcoding and acidification was recorded by pH monitoring.

The raw milk samples investigated in this study showed high bacterial diversity. The microorganisms present in raw milk can have different origins including the animal’s teat, the dairy equipment, and the animal’s surroundings. Although hygiene practices induce a strong decrease in microbiological load, leading to a stable low microbial load of approximately 10^3^-10^4^ CFU/mL (37), raw milk microbiota is still characterized by high diversity (38–41). The majority of raw milk samples were enriched in *Pseudomonas*, which are psychrotrophic bacteria. After dispersal, i.e. after transfer of microorganisms from the environment into the milk (42), milk samples were stored at 4 °C for up to 72 hours until fermentation. This step could have led to the enrichment of milk in psychrotrophic bacteria as previously described (39, 41). After the first fermentation step, a drastic reduction in bacterial diversity was observed. This reduction likely resulted from the strong selective pressure that applies during fermentation and only allows certain microorganisms such as LAB to persist (42).

Although the lag time for the first fermentation step was long (approximately 14 hours), which is in agreement with previous reports (43), the successful spontaneous fermentation of all raw milk samples implies that they were all initially colonized by microorganisms able to acidify milk. In line with this, metabarcoding analyses revealed that all were colonized by LAB, which can be considered as the main contributors to milk fermentation. Thus, even though LAB were present in very low relative abundance in raw milk, they were able to assure milk acidification, suggesting that adaptation to the milk matrix is more determinant than the relative abundance of bacteria initially present in raw milk. Besides LAB, several, but not all, fermented milk samples were colonized by other bacteria including species of the phylum *Pseudomonadota*. The impact of these bacteria on cheese is poorly described in the literature. Gram-negative bacteria such as *Escherichia coli* exclusively considered as spoilage bacteria for years as members of this phylum are unwanted because of their animal intestine origin. However, some Gram-negative bacteria such as *Psychrobacter celer* or *Hafnia alvei* could have a beneficial impact on cheese by conferring aroma development and protection against pathogens (44, 45). Top-down community engineering through raw milk spontaneous fermentation could also be an interesting source of beneficial bacteria other than LAB, including Gram-negative bacteria. It can be noted, however, that the direct use of such communities for applications could pose problems for health reasons. Indeed, very few Gram-negative bacteria are authorized for use as starter cultures. Therefore, it would probably be more favourable to use those communities that contain poorly understood or undesirable microorganisms in an integrated design approach (11): after the acquisition of communities by top-down design, a bottom-up design approach could be applied to build synthetic communities excluding these undesirable microorganisms.

Although richness decreased in our experimental engineering setup, it was still high compared to the communities previously described and obtained by advanced backslopping (22). Indeed, Swiss hard cheese starter cultures were colonized by two LAB species: *Streptococcus thermophilus* and *Lactobacillus delbrueckii* subsp. *lactis.* By contrast, most of the fermented milk samples obtained in our study were colonized by approximately 13 different ASV on average, including different species. This suggests that early backslopping, starting from raw milk, could be better suited to capturing microbial diversity and could be interesting for the design of starter cultures.

In this study, 3 groups based on the change in lag time, the MAR and the final pH of the acidification curves were distinguished. GP1 showed a reproducible profile from F2 to F6, GP2 was more variable, and GP3 showed higher variations with intersecting curves. Metabarcoding analyses revealed that changes in community structures mirrored the acidification patterns: GP1 showed stable communities from F2 to F6, mainly dominated by one LAB genus (*Lactococcus* or *Streptococcus*), GP2 showed moderate variations (with more diversified communities) and GP3 showed the highest variations (with a dominance of *Lactococcus* in the first steps and a dominance of *Lactobacillus sensu lato* in the last steps). Milk is a medium with high concentrations of nutrients. It was previously shown that serial propagation of communities in nutrient-rich media leads to communities with low diversity and high shaping (i.e. low stability) (46). In nutrient-rich media, bacteria greatly alter the chemical environment, causing more negative interactions between species. Strong species interactions could explain the reduction in bacterial diversity during milk fermentation performed in our study. However, besides the lineages characterized by low stability (in GP2 and GP3), some lineages were stabilized in a few propagation steps. The study of Ratzke *et al.* was conducted following a bottom-up design approach where 15 different soil bacteria were mixed to build synthetic communities (46). The serial propagations conducted in our study involved 26 milk samples, each colonized by an average of 60 ASV. The higher diversity sampled in our study could have allowed us to highlight cases where community stabilization is possible and gives rise to the GP1 communities. The GP1 communities were less diversified than GP2 and GP3 communities. This suggests that conservative propagation during backslopping is facilitated when communities have a relatively low diversity, which would be consistent with data obtained in previous studies (22, 43).

Community lineages with a stable final pH also presented reproducible metabarcoding profiles. Furthermore, although GP1 communities are stabilized, they not only displayed interesting structural diversity (as shown by NMDS) but also functional inter-lineage diversity. Indeed, the final pH of GP1 communities differed significantly between the lineages in this group. Thus, in contrast to previous work (21, 22), which showed a high degree of functional redundancy between communities obtained by advanced backslopping, here we show that it is possible to obtain community lineages exhibiting dissimilar community structures at the interspecific level, and presenting different functions. In cheesemaking, the pH varies strongly depending on the technology (47). The possibility of obtaining communities with different acidification properties opens up very interesting prospects for innovation in the field of dairy processing. More precisely, the starters used in the dairy sector are cultivated in high amounts and are added at different stages of the process. However, these starter cultures contain a low species diversity and are usually used by a lot of cheesemakers (48). Our results show that early backslopping could be a valuable tool for selecting communities that can be used as starter cultures to restore microbial diversity in dairy products.

Our results showed that the dominance of LAB over *Pseudomonadota* led to both a lower MAR and a lower final pH. More precisely, these parameters depended on the dominant LAB genus *Lactobacillus sensu lato*, *Lactococcus* or *Streptococcus*. The highest lag times were associated with *Lactobacillus sensu lato*, in comparison with the other LAB or the *Pseudomonadota*. In GP3, the ecological trajectories of the lineages were the longest and tended to be parallel, reflecting the microbial community succession in this group. *Lactococcus* was dominant in the first fermentation steps, and then decreased while *Lactobacillus* increased and became dominant in the final steps. Interestingly, the level of dynamics was associated to the presence of *Lactobacillus sensu lato*: the higher the relative abundance of *Lactobacillus sensu lato* in the lineages, the higher the dynamics. This suggests that *Lactobacillus sensu lato* has a strong influence on the balance between community shaping and propagation. This role of lactobacilli on community dynamics could be due to their organic acid production and to their resistance to acid stress. *Lactobacillus sensu lato* can actually produce more acid and can be more resistant to acid stress than *Lactococcus* (7, 49, 50). In addition, Herreros *et al.* explained that lactobacilli metabolize lactose more slowly than lactococci (51). These differences in acid production and tolerance could explain the change in pH, and the exclusion of *Lactococcus* to the benefit of *Lactobacillus,* in the GP3 lineages. Alternatively, the major changes observed in GP3 could be due the presence of bacteriophages in the communities. Phages are known to target lactococci (52) and could induce a decrease of these LAB in the community, which could have allowed the *Lactobacillus* to become dominant in the GP3 communities. Other studies have indeed shown that bacteriophages are responsible for population dynamics during backslopping, albeit at the intraspecific level (21, 22, 43, 53).

## Conclusion

In conclusion, this work has shown the wide diversity of community behaviour in terms of dynamics, diversity and trajectories, resulting in significant functional impacts in the framework of a top-down community engineering approach. The use of dairy backslopping at early stages is a powerful model to investigate the impact of microbial diversity and interactions on the balance between community shaping and propagation. Furthermore, this work on the top-down engineering of complex microbial communities offers very interesting prospects in all areas where expectations are high in terms of restoring microbial diversity in ecosystems.

## Data availability

The datasets presented in this study can be found in the data repository Recherche Data Gouv at https://doi.org/10.57745/0XZAQN

## Acknowledgments

Funding was provided by the Association Nationale de la Recherche et de la Technologie (CIFRE 2020/1026 agreement).

The authors thank Annelore Elfassy, Amandine Martin, Myriam Schivi and Sylvie Wolff from LIBio, Béatrice Desserre, Isabelle Verdier-Metz, Céline Delbès and François Allegretti from UMRF, for their advice and technical help.

Conceptualization: CG, FB, CM, CCh, CCa; Data curation, formal analysis, resources and software: CG, AD, FB, ST; Funding acquisition: FB, CCh, AMRJ; Investigation and writing - original draft: CG; Methodology: CG, FB, CCh, CCa, AD, ST; Project administration: CG, FB, CCh; Supervision: FB, CCh, CM, CCa, AMRJ; Validation: FB, CM; Visualisation: CG, FB, AD; Writing - review & editing: CG, FB, CM, AD, CCh, CCa.

## Supplementary figure legends

**Figure S1. Acidification kinetics of all raw milk samples during serial fermentation.** The colours indicate the fermentation step, from light pink (F1) to dark red (F6).

**Figure S2. Acidification parameters according to the fermentation step (F1-F6) for all samples.** (A) Lag time and (B) MAR. Different superscript letters indicate a significant difference (*p* < 0.05; pairwise comparisons using the Wilcoxon rank sum exact test with the Bonferroni adjustment method for the *p* value).

**Figure S3. Alpha diversity (richness) of the raw milk samples.** (A) According to the milk sample and (B) according to the acidification group. The same superscript letters indicate no significant difference (*p* > 0.05; one-way ANOVA).

**Figure S4. Alpha diversity of all communities according to the fermentation step (T0-F6).** (A) Richness, (B) Shannon index. Different superscript letters indicate a significant difference (*p* < 0.05; Pairwise comparisons using the Wilcoxon rank sum exact test with the Bonferroni adjustment method for the *p* value).

**Figure S5. Bacterial community composition at the phylum level for each raw milk sample and fermentation step.** T0: raw milk samples, F1 to F6: fermented milk samples.

**Figure S6. Alpha diversity analysis.** (A, B) Richness for each lineage and overall (F2-F6) and (C) sum of richness variations between adjacent steps, for each group GP1, GP2 and GP3. (D, E) Shannon indices for each lineage and overall (F2-F6) and (F) sum of Shannon indices for each group. Different superscript letters indicate a significant difference (*p* < 0.05; Pairwise comparisons using the Wilcoxon rank sum exact test with the Bonferroni adjustment method for the *p* value).

